# Capturing the developing brain in motion: A practical tutorial for recording mobile EEG in naturally moving children

**DOI:** 10.64898/2026.07.22.739990

**Authors:** Annika Werwach, Sarah D. Power, Markus Werkle-Bergner, Ulman Lindenberger, Sarah Jessen

## Abstract

This tutorial seeks to facilitate the use of mobile electroencephalography (EEG) in young children. Mobile EEG allows researchers to investigate neural correlates of behavioral and cognitive processes in ecologically valid settings. While mobile EEG has been widely adopted in adult research, studies applying it to freely moving children remain scarce. However, investigating neural processes during active behavior and in interaction with movement is indispensable for advancing our understanding of neural correlates of cognitive development in early childhood. Here, we provide a practical tutorial on the collection and preprocessing of mobile EEG data from children. Drawing on experience and data from a large-scale study with toddlers, we summarize key methodological considerations, discuss common challenges and practical recommendations, and present a preprocessing pipeline developed for developmental mobile EEG data. In addition, we provide example EEG datasets from naturally moving 18-20-month-old toddlers. We argue that incorporating mobile EEG into developmental cognitive neuroscience allows researchers to (i) investigate cognitive development in naturalistic environments, (ii) examine associations among bodily movement, neural activity, and behavior, and (iii) study samples that are difficult to reach with stationary research.

**Highlights:**

- Mobile EEG enables investigation of neural processes during active behavior
- Mobile EEG in moving freely children comes with unique methodological challenges
- We provide practical recommendations for developmental mobile EEG studies
- A dataset and preprocessing pipeline from a toddler mobile EEG study are provided

## 1. Introduction

Children develop through active interactions with their social and physical environment. Already during infancy, they actively seek out contexts for learning (Begus and Southgate, 2012; Jirout et al.; 2024, Ke et al., 2025), and curiosity-driven exploration positively relates to cognition across childhood (de Boer et al., 2026; Jirout et al., 2024; Muentener et al., 2018; Ziv et al., 2026). Movement and cognition are tightly linked from early on (e.g., Robertson et al., 2001), with both active exploration (Chung et al., 2024; Lozada and Carro, 2016) and general motor development and movement experience (Chung et al., 2024; Colomer et al., 2023; Pacheco et al., 2025; Schwarzer and Jovanovic, 2024) facilitating cognitive development throughout childhood.

However, despite the central role of active exploration, interaction, and movement in development, most studies examine children’s cognitive processes and their development in highly controlled laboratory settings (see Dahl, 2017; van Atteveldt et al., 2018). Such settings have provided crucial insights into cognitive development as they allow for precise experimental control and the isolation of specific cognitive mechanisms. However, they usually rely on isolated and standardized stimuli and strongly constrain children’s natural behavior (see Shamay-Tsoory and Mendelsohn, 2019). As a consequence, important aspects of cognitive development, such as active exploration, spontaneous movement, and interaction with complex environments, are underrepresented (Cantlon, 2020; Dopierala and Emberson, 2023; Janssen et al., 2021; Wass and Goupil, 2022). Hence, laboratory-based studies need to be complemented with more naturalistic approaches, examining children’s behavior and cognitive processes in real-world environments (see Dahl, 2017; Janssen et al., 2021; van Atteveldt et al., 2018; Wass and Goupil, 2022).

Behavioral research has started to address the gap between highly controlled laboratory studies and children’s real-world experiences by studying children’s behavior and cognitive processes in more naturalistic settings (e.g., Dahl, 2015; van Liempd et al., 2025). For example, Pathman and colleagues (2023) examined free recall in 4- to 10-year-olds based on their experiences in a summer camp at a zoo, showing how children retrieve information in naturalistic environments. While these behavioral studies provide important insights into cognitive processes and their development in real-world contexts, the neural mechanisms underlying these processes remain elusive, in large part due to the additional challenges of recording neuroimaging data in freely moving children. Importantly, neural measures can provide complementary insights into the mechanisms through which children perceive, process, and learn from their environment that cannot be inferred from behavior alone.

It is increasingly recognized that neuroimaging studies in naturalistic settings, which allow participants to move and interact freely within real-world environments, are essential for a complete understanding of cognition and behavior (see Hille et al., 2024; Makeig et al., 2009; Maselli et al., 2023; Stangl et al., 2023; van Atteveldt et al., 2018). While adult neuroscience has successfully demonstrated the usability of portable neuroimaging techniques such as mobile EEG to investigate the neural mechanisms underlying movement and motor-related cognitive mechanisms, (Gwin et al., 2011; Jain et al., 2013; Klapprott and Debener, 2024; Storzer et al., 2016), as well as general cognitive processes during active behavior (Klapprott and Debener, 2024; Papin et al. 2024; Piñeyro Salvidegoitia et al., 2019; Scanlon et al., 2017), mobile neuroimaging studies allowing for free movement in early and middle childhood have remained scarce. Most mobile EEG studies with children have been conducted in classroom settings or in children’s homes (Chin et al., 2025; Dickinson et al., 2025; Troller-Renfree et al., 2021), demonstrating the feasibility of EEG recordings outside of laboratory environments in children (see Box 1 for further examples of recent studies). However, these studies typically still impose substantial restrictions on participants’ behavior and movement (e.g., requiring children to remain seated and minimize movement). In addition, stimuli are usually highly standardized, for example presented on computer screens or as structured classroom lessons. As a result, mobile EEG studies in childhood that allow for active behavior and free movement in naturalistic contexts remain scarce.

Such active and unconstrained behavior however is essential to study cognition in childhood (see Amadó et al., 2024; Byrge et al., 2014; Dopierala and Emberson, 2023; Janssen et al., 2021; Mathewson et al., 2024) since cognitive development emerges through ongoing interactions between perception, action, and children’s social and physical environment (see Kontra et al., 2012; Marmeleira and Duarte Santos, 2019; Schwarzer and Jovanovic, 2024). Studying these processes under naturalistic conditions can thus complement findings from laboratory studies and facilitate the generalization of lab-based results to ecologically valid contexts. Beyond its potential for studying these broader research questions, mobile EEG may also hold specific practical advantages for developmental cognitive research. This may be particularly relevant for young toddlers, whose development is characterized by active exploration and movement (Adolph and Hoch, 2019; Jirout et al., 2024; Ke et al., 2025), but who are also among the most difficult populations to study using traditional stationary EEG paradigms because of the associated behavioral and movement constraints. Similar advantages may also apply to other populations that have difficulty complying with stationary paradigms, such as children with neurodevelopmental disorders. Explicitly allowing children to move and express their natural behavioral tendencies during mobile EEG data acquisition can thus not only address naturalistic research questions but also allow the inclusion of populations that are difficult to study in traditional stationary research.

### Box 1: Recent mobile EEG studies in childhood and adolescence

Chin et al., 2025: Strategies to collect resting-state and visual evoked potential mobile EEG data from 24-month-old children in rural Ethiopia

Dickinson et al., 2025: Mobile EEG data collected in home environments from 4-year-olds children maintains data quality and neural signal integrity comparable to laboratory EEG recordings

Nicklas et al., 2026: Cross-domain integration (motor, cognitive, sensory) in adolescents (13-23 years) with ASD differs from typically developing controls

Troller-Renfree et al., 2021: Methodological and analytical guidance for collecting resting-state mobile EEG data from 12-month-old infants in home environments

Xu et al., 2022: Mobile EEG can be used in 6-7-year-old elementary school children to examine neural correlates of attention during classroom lessons

While mobile EEG approaches therefore hold great potential, collecting EEG data from naturally moving, young children also comes with substantial methodological challenges. These include unique movement-related artifacts, participant compliance, task-design considerations, and the suitability of available EEG equipment and preprocessing pipelines. In the present tutorial, we show how these challenges can be addressed. Our focus is on the collection and preprocessing of mobile EEG data in early childhood (see Box 2 for the definition of mobile EEG used in the context of this tutorial). Drawing on experience from a large-scale, longitudinal mobile EEG study with freely moving toddlers, we summarize key methodological considerations, common challenges and problems, and practical solutions. In addition, we present an example dataset collected from 10 freely moving toddlers aged 18-20 months and an accompanying preprocessing script, report statistics on data quality and conduct a comparison to an established developmental EEG preprocessing pipeline. By demonstrating the feasibility of recording neural activity during active behavior in this particularly challenging age group, we aim to lower the barriers to implement mobile EEG in research in early and middle childhood and facilitate the investigation of neural processes during naturalistic behavior in early childhood.

### Box 2: What do we mean by mobile EEG?

In the context of this tutorial, we define *mobile EEG* as electrophysiological recordings acquired while participants are moving freely within their environment (e.g., crawling, walking, turning, interacting with objects), without substantial movement constraints imposed by the experimental setup. In contrast, *stationary EEG* refers to recordings during which participants’ movement is strongly restricted, usually requiring seated and relatively motionless behavior.

Importantly, mobility in EEG studies can be defined along two dimensions: the degree of participants’ own movement and the portability of the recording system. Consequently, EEG studies lie on a continuum with respect to mobility. For example, some studies allow limited participant movement (e.g., treadmill walking, children seated on parent’s lap), while the recording hardware remains fixed (e.g., Gramann et al., 2010, Liao et al., 2015). Other studies use portable or wearable amplifiers, but participants are still required to be seated during tasks (e.g., Bevilacqua et al., 2019, Ko et al., 2017).

In this tutorial, we primarily focus on settings in which both participants and the recording system are fully mobile and unconstrained with regards to movement. However, many methodological considerations discussed here will also apply to partially mobile EEG settings.

## 2. Example dataset

The example dataset associated with this tutorial contains data from *n* = 10 naturally moving toddlers aged 18-20 months (5 girls). The data were recorded during the encoding session of a longitudinal study on episodic memory development at the Max Planck Institute for Human Development in Berlin, Germany (Power et al., in prep.). Toddlers performed a hide-and-seek task in which they searched for a toy in different boxes until they found the toy. The study followed American Psychological Association standards in accordance with the declaration of Helsinki from 1964 and was approved by the local Ethics Committee of the Max Planck Institute for Human Development (reference numbers: LIP-2022-03/ECHO and LIP-2023-03/ECHO). Parents provided written informed consent prior to participation. In total, *n* = 303 mobile EEG datasets were collected in this study across three different sessions in a longitudinal design. In the encoding session, *n* = 105 EEG datasets were collected.

EEG data was recorded separately for each trial, together with videos of toddlers’ movement, with a trial starting with the child entering the experimental space and ending with the child leaving it after successfully locating the toy. The mean number of trials in the example datasets is 6.4 (*SD* = 2.37, *min* = 4, *max* = 11), which is representative of the full sample for this first test session (*n* = 105; *M* = 6.4, *SD* = 2.43, *min* = 4, *max* = 13). EEG and videos were synchronized via LSL (Lab Streaming Layer; Kothe, 2014) and saved in .xdf files.

Example data and preprocessing scripts can be found here: osf.io/x5tp8/

## 3. Practical challenges and recommendations for mobile EEG in childhood

### 3.1. Mobile EEG equipment and setup

The growing number of mobile EEG studies in adults has led to the development of an increasing number of mobile EEG systems. Both consumer- and research-grade systems are available and have been used in research (for a general overview and discussion, see e.g. Gramann et al., 2011; Lau-Zhu et al., 2019; Mathewson et al., 2024; Niso et al., 2023). These systems differ with respect to several aspects, including electrode type, amplifier portability, or wireless transmission technology; for a comprehensive overview, see Niso and colleagues, (2023). In this tutorial, we focus primarily on research-grade systems and highlight considerations that may be particularly relevant for studies with children.

Small, lightweight, and comfortable systems are generally recommended in mobile setups with adults (see Makeig et al., 2009), and even more preferable in early childhood studies. Comfort, ease of use, and a short preparation time can reduce attrition rates, especially in young children. Because gel-based systems often require long preparation, some research groups thus favor dry electrode systems for child samples (e.g., Troller-Renfree et al., 2021). However, dry systems can be particularly susceptible to movement-related artifacts and high impedance levels (Luu et al., 2024; Mathewson et al., 2017), potentially leading to considerable noise and data loss in studies involving movement. Since EEG recordings from children usually contain a high rate of artifacts even in stationary settings, this limitation becomes especially relevant for developmental mobile EEG studies. For the example data of this tutorial, we therefore opted for a gel-based system. To reduce preparation time, we recommend preparing the cap with gel before capping the child (pre-gelling, see Turk et al., 2022). Note that this approach requires specific training to avoid gel bridges between electrodes due to child movement during cap application.

Mobile EEG systems come with wired or wireless configurations. When using wired systems with young children, cables should be secured and positioned outside of the child’s reach and field of view to minimize the risk of children pulling on the cables. Amplifier placement also needs to be considered carefully in mobile EEG studies. Some systems use a small amplifier that is not connected to the cap itself, requiring researchers to find a suitable placement of the amplifier on the participant’s body. In our experience, placing the amplifier in a small backpack carried by the child keeps amplifier and cables outside of children’s reach and field of view. However, such setups require an additional item of equipment, which may be challenging depending on the age group (see below, section 3.3.1), and can introduce additional artifacts to the data caused by cable movement. In other systems, the amplifier is integrated within the cap, reducing cable-related artifacts and necessary equipment items. When using these setups with young children, however, researchers need to ensure that participants are not hurt when falling onto the cap, which is more likely in mobile studies with young children compared to adults. In addition, caps with integrated amplifiers may be uncomfortable for infants when they lay down.

Many mobile EEG setups also include auxiliary channels such as EOG electrodes or motion sensors to capture additional movement variables. In our experience, additional electrodes or sensors can lead to irritation and resistance in young children or even an increased drop-out rate, particularly when placed in the face. For this reason, we generally recommend limiting additional equipment to what is essential for addressing the research question.

### 3.2. Experimental design

More so than adults, young children tend to only actively participate in studies when they are comfortable and intrinsically motivated. Thus, a major challenge for all EEG studies with young children is the design of age-appropriate, child-friendly, and fun tasks requiring only minimal and potentially non-verbal instructions. In this regard, mobile EEG tasks may even have an advantage over stationary set-ups: Since young children usually enjoy moving around and exploring their environment, mobile tasks are generally more likely to align with children’s behavioral tendencies than stationary tasks.

Depending on the age group of interest, mobile EEG tasks need to be adapted to the motor skills of this specific age group. For example, a search task is only possible once toddlers are able to crawl or walk safely. Other movement behaviors, such as running, cycling, jumping, or performing sports, can only be studied in older children. Additionally, as is also the case for stationary research with children, the duration of the task should be adapted to the attentional capabilities of the chosen age group (see Turk et al., 2022, for age-specific recommendations). Ideally, paradigms should be designed in a way that data are usable even if children are not able to complete the entire duration of the task. Additionally, because children may deviate from instructions during the experiment, experimental protocols should include ways to handle extensive movement variability across time and space.

Consideration should also be given to the tradeoff between naturalistic movement and the signal-to-noise ratio in the EEG signal. Young children can behave and move more unpredictably than adults, such as spontaneously starting to run or roll on the floor, adding more variable and stronger noise to the signal. The degrees of freedom in children’s movement can be influenced by the experimental design to some extent, such as constraining the space in which they can move. To push the boundaries of what is possible in terms of data analysis, the dataset provided alongside this paper provides an example of relatively unconstrained movement, including running, crawling, hopping, being lifted up and down etc.

### 3.3. Data collection

Data collection procedures for mobile EEG in children strongly influence attrition rates, children’s ability to focus on the task, and quality of EEG recordings. In the following, we focus on strategies to increase participant engagement and compliance during application of the EEG system and recording procedures as well as data quality monitoring that should be considered in the context of mobile EEG studies with young children.

#### 3.3.1. Participant engagement and compliance

Compliance strategies for EEG data collection in different age groups of children, such as short video clips, a designated experimenter for distraction and entertainment, or interesting toys and snacks, have been discussed elsewhere (e.g., Bell and Cuevas, 2012; Chin et al., 2025; Norton et al., 2022; Troller-Renfree et al., 2021; Turk et al., 2022). The collection of mobile EEG data often comes with additional equipment besides the EEG caps, such as storage solutions for amplifiers, movement sensors, or auxiliary electrodes. In our experience, the use of child-friendly terminology when explaining equipment items, allowing children to touch parts of the equipment when possible, and also a certain flexibility (e.g., using child’s own backpack instead of the study back-pack) can increase acceptance. If children are scared of the equipment, actively involving parents and having adult-size equipment for parents to wear themselves often makes children more comfortable. Given that, depending on the age group, some children may refuse to wear all equipment items, we recommend defining both necessary items for study inclusion as well as optional items that are not necessary for study inclusion. A possible approach is to collect data from a large behavioral cohort and one or multiple sub-cohorts of children who accepted the EEG equipment and potential additional sensors. Regardless of inclusion procedure, researchers should keep in mind that developmental EEG studies are generally subject to selection bias, as children who tolerate the EEG equipment may differ systematically from those who do not. This bias may be particularly relevant for mobile EEG studies involving additional equipment items and sensors and should be considered when interpreting findings.

#### 3.3.2. Recording procedures and data quality monitoring

For recordings outside of laboratories, experimenters need a site where both the EEG cap administration as well as de-administration can take place, including access to water, spaces for the children to sit, and storage of consumables such as Q-tips or tissues, gel, etc. This place should be designed in a child-friendly way and include toys or other possibilities to distract the child. Additionally, when gel-based systems are used, there should be a place for washing the child’s hair after the experiment. In our experience, a change in location and context between cap and equipment administration and data collection can serve as a distractor for children, increasing acceptance of the cap and task.

Impedance assessments are a way to monitor conductance and signal quality before the start of the experiment. In mobile paradigms, it is more likely that caps move or children sweat and pull on electrodes. For this reason, it may be sensible to add additional online measures of data quality and to examine impedances again at the end of the experiment.

When collecting data in naturalistic tasks, the recording of triggers during the experiment can be challenging. In contrast to traditional EEG recordings, where the presentation of stimuli and the registration of corresponding triggers is timed with millisecond precision, it is much more difficult to obtain precise event triggers in naturalistic, mobile experiments. The choice of an appropriate trigger recording procedure strongly depends on the research questions and analysis aims: If analyses allow triggers to be relatively imprecise in the time domain (e.g., induced oscillatory power changes), the recording of manual triggers during the experiment may be sufficient. However, if research questions focus on ERPs, other solutions need to be found to precisely time triggers to the targeted actions, such as extracting event timings from video recordings, or using motion tracking via motion sensors. In both cases, the video or motion tracking streams should be precisely synchronized to the EEG signal, for example via the software Lab Streaming Layer (LSL; Kothe, 2014). Adult studies have developed additional methods to register events during mobile EEG recordings, such as time-locking the EEG data to blink-, saccade-, or gait-related events (see Wunderlich and Gramann, 2021), which may also have potential for the use in children.

### 3.4. Preprocessing and movement artifacts

Both mobile EEG and developmental EEG are known to be prone to a high rate of artifacts. For this reason, when using mobile EEG with children, effective preprocessing steps to increase signal-to-noise ratio are crucial. Standardized preprocessing pipelines are available for both infant and child recordings, such as the MADE (The Maryland analysis of developmental EEG pipeline, Debnath et al., 2020), HAPPE (The Harvard Automated Processing Pipeline for Electroencephalography, Gabard-Durnam et al., 2018), and APICE pipelines (Automated Pipeline for Infants Continuous EEG, Fló et al., 2022), as well as for mobile EEG data (BeMoBIL pipeline, Klug et al., 2022). However, there are currently no pipelines tailored to both, the specific characteristics of mobile EEG and the specifics of EEG data recorded from young children. Hence, we provide an example pipeline that was used to preprocess mobile EEG data from 18-20-month-old toddlers.

### 3.5. Summary and key recommendations for mobile EEG studies in children

Mobile EEG in children requires adapting all stages of the research process to children’s characteristics and the specific requirements of naturalistic recordings. When planning and designing mobile EEG studies with children, we recommend to:

1. develop simple, child-friendly, and safe recording setups that can be prepared in advance and minimize the amount of equipment worn by the child;
2. adapt experimental designs and tasks to children’s motor and attentional capabilities;
3. allow for variability in behavior and recording duration;
4. implement careful data-quality monitoring and preprocessing strategies to handle movement-related artifacts;
5. utilize the benefits of mobile EEG for early and middle childhood populations to increase motivation and compliance and allow the inclusion of subgroups currently underrepresented in developmental EEG studies.

## 4. Developmental mobile EEG preprocessing pipeline: Tutorial and recommendations

### 4.1. Pipeline overview

This preprocessing pipeline was developed specifically for mobile EEG data collected in 18-20-month-old toddlers, in the context of a study on memory development in the second year of life. It aims to combine experiences and recommendations from both developmental and mobile EEG preprocessing pipelines in order to address the unique characteristics and challenges of mobile EEG data collected from children. Importantly, we do not aim to provide a standardized pipeline. Instead, we share our experiences preprocessing EEG data collected in a mobile paradigm with relatively unconstrained movement, which may guide others who plan to conduct mobile EEG studies with young children. Figure 1 provides a flowchart of the pipeline. Please note that parameters where optimized for 18-20-month-old children performing a hide-and-seek task and may require adaptation for other age groups and experimental paradigms. We report descriptive statistics on the number of excluded data segments and channels, artefactual ICs, and segments containing artifacts after preprocessing based on the full sample processed with this pipeline (*n* = 303). The provided example datasets (*n* = 10), which are a subset of this full sample, can be used to inspect and reproduce the preprocessing steps.

**Figure 1.**
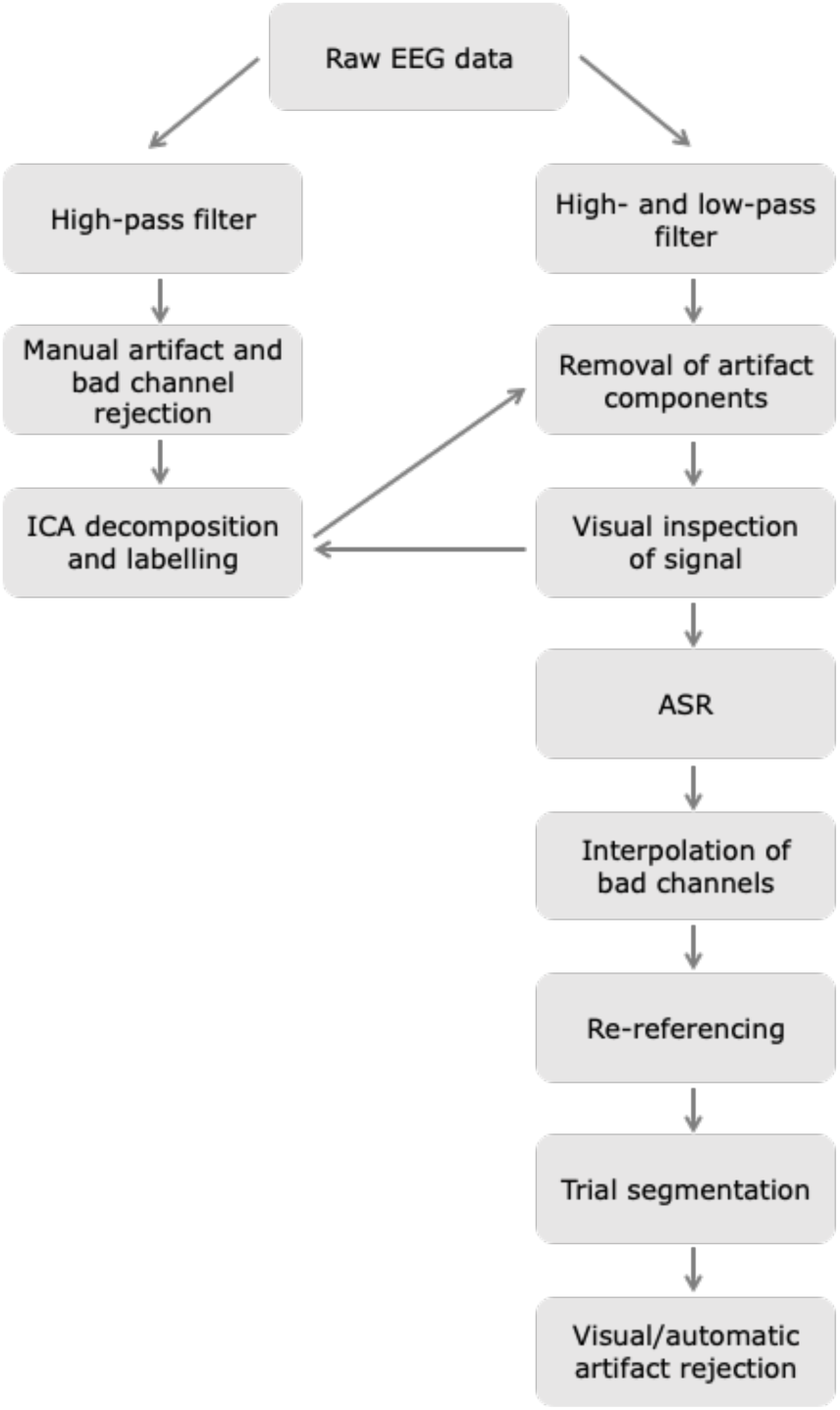
Visualization of preprocessing pipeline developed for mobile EEG recordings from naturally moving toddlers.

Importantly, this pipeline mainly comprises preprocessing techniques that are frequently applied in most EEG settings. We thus demonstrate that even if the raw data are strongly contaminated by movement- and motion-related artifacts, commonly used preprocessing steps can result in EEG data that are usable for further analyses. See Figure 2 for example EEG segments before and after preprocessing.

**Figure 2.**
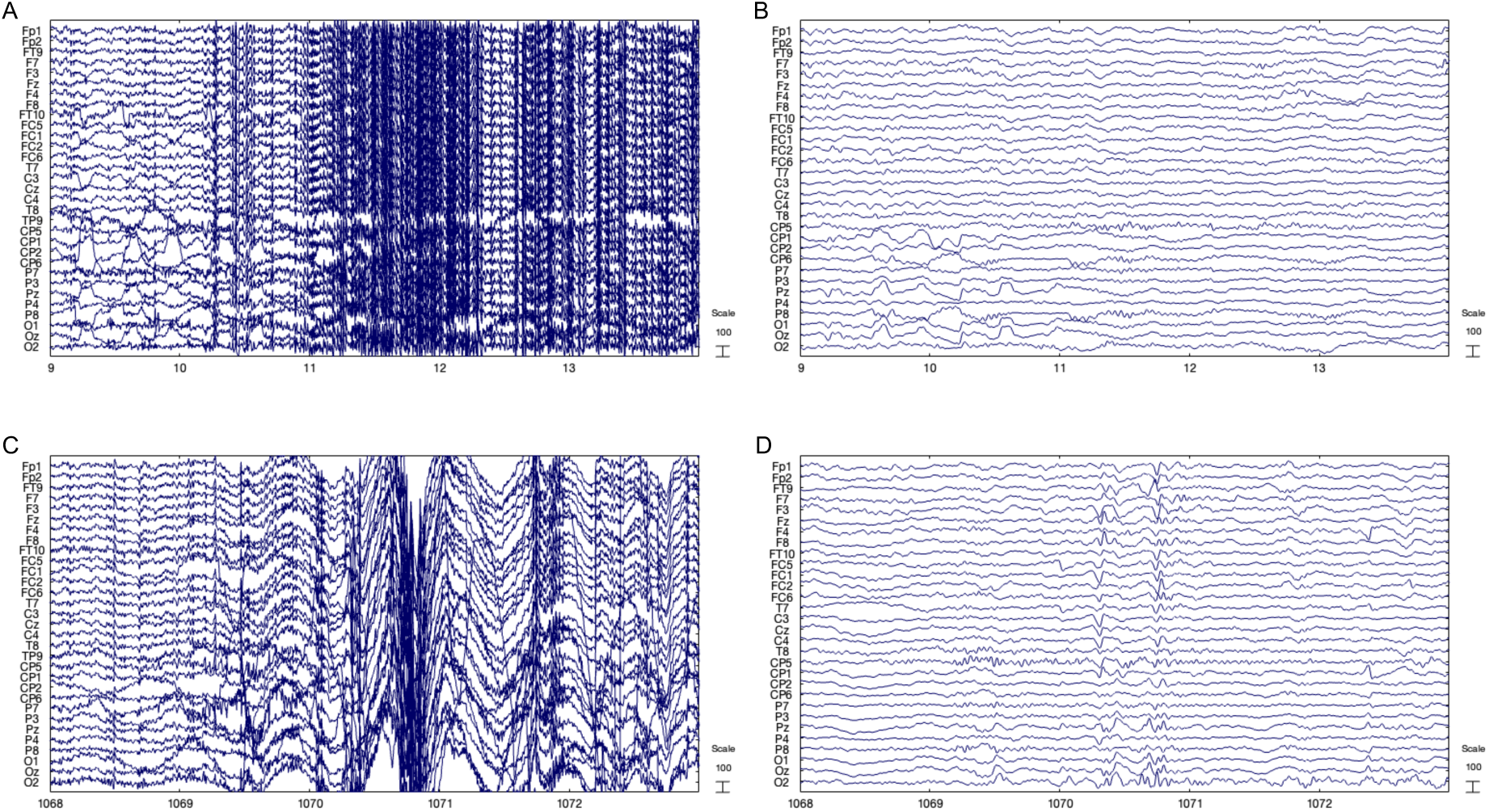
Example EEG segments from the first example dataset (sub-001) before (A, C) and after (B, D) preprocessing with the custom pipeline. Panels A and B show a 5s segment from 9–14 s of the continuous recording, while panels C and D show a second 5s segment from 1068–1073 s. The raw data were high-pass filtered at 1.25 Hz prior to preprocessing.

All preprocessing steps were done using MATLAB version R2024b (The MathWorks Inc., 2024) and the EEGLAB (Delorme and Makeig, 2004) and Fieldtrip toolboxes (Oostenveld et al., 2011).

### 4.2. Step-by-step pipeline

#### 4.2.1. Step 1: High-pass filter

In the first step, EEG data were high-pass filtered. The sole use of a high-pass filter without a low-pass filter is recommended to improve the independent component (IC) decomposition for mobile EEG data (Gorjan et al., 2022; Klug et al., 2022; Klug and Gramann, 2021). For the same reason, it is also recommended in both mobile and developmental pipelines to use a high-pass filter cut-off between 1-2 Hz (Gabard-Durnam et al., 2018; Gorjan et al., 2022; Klug et al., 2022; Winkler et al., 2015). We therefore used a cut-off of 1.25 Hz (filter order: 3300, passband edge: 1.5 Hz, stopband edge: 1.0 Hz) with a zero-phase Hamming-windowed sinc FIR filter (see Debnath et al., 2020; Fló et al., 2022; Klug et al., 2022).

#### 4.2.2. Step 2: Artifact rejection for Independent Component Analysis (ICA)

Subsequently, the EEG signal was manually scanned to remove 1s-segments containing severe artifacts as well as bad channels, as preparation for the Independent Component Analysis (ICA; Makeig et al., 1996). We decided against using automatic artifact and bad channel detection methods that were trained on stationary data because of the unique nature of the high-amplitude artifacts included in the signal and the general strong contamination of the signal with artifacts. Importantly, only the most severe artifacts and channels with almost no usable signal were removed from the data in order to keep enough data for a usable IC decomposition. On average, 206.2 1s-segments were rejected per dataset, with 1181.2s of data being retained on average (average length of datasets before artifact rejection: 1387.4s).

#### 4.2.3. Step 3: Independent Component Analysis (ICA)

Based on topography and waveform, ICs were manually classified as being likely to represent neural activity, eye movements, muscle or movement artifacts. Again, we decided against automatic classification algorithms (e.g., ICLabel, Pion-Tonachini et al., 2019; adjusted-ADJUST, Leach et al., 2020; or MARA, Winkler et al., 2014, Winkler et al., 2011), which have been trained on stationary data from adults or children. The reason for this was the high number of non-stereotypical artefactual ICs derived from the data. We rejected an average of 8.33 artifact ICs out of max. 31 total ICs (*SD* = 2.85, min.: 1, max.: 20). Artefactual ICs included the following categories: noise or muscle artifacts from the reference electrode; single- or multi-electrode muscle components (often from outer electrode ring); single-electrode noise; mechanical artifacts affecting all or a subset of electrodes at the same time; eye electrode noise due to pulling or excessive face movements; blinks; horizontal eye movements; other eye movements.

#### 4.2.4. Step 4: High- and low-pass filter

Following IC classification, the raw EEG signal was loaded again and high- and low-pass filtered. We decided for separate high- and low-pass filters and a transition bandwidth of 0.5 Hz for the high-pass and of 5 Hz for the low-pass filter in order to minimize artifacts introduced by steep filter roll-offs while maintaining enough signal at the lower frequency end (see Widmann et al., 2015). The chosen cut-off of the high-pass filter was 0.5 Hz (filter order: 3300, passband edge: 0.8 Hz, stopband edge: 0.3), and the cut-off of the low-pass was 30 Hz (filter order: 330, passband edge: 27.5 Hz, stopband edge: 32.5 Hz), both using a zero-phase Hamming-windowed sinc FIR filter. The low-pass filter cutoff was set at 30 Hz to remove muscle-related activity in the EEG signal, which usually extends across frequencies from 30 Hz on (see Muthukumaraswamy, 2013).

#### 4.2.5. Step 5: Removal of artifact ICs

Artifact ICs identified before were then removed from the high- and low-pass-filtered signal. Crucially, the resulting signal was again visually inspected, and if high-amplitude, stationary artifacts persisted, the ICs identified based on the high-pass filtered data were inspected again and additional components that corresponded to the residual artifacts were identified. This iterative process was repeated until no artifacts that may be removed via the ICA procedure could be identified in the signal.

#### 4.2.6. Step 6: Artifact Subspace Reconstruction (ASR)

After the removal of artifactual ICs, Artifact Subspace Reconstruction (ASR; Kothe and Makeig, 2013; Kothe and Jung, 2016) was performed using a cut-off parameter of 10 to correct for remaining severe artifacts in the data. Because the present dataset contains very few (4–13) and long trials (usually multiple minutes), traditional artifact rejection procedures rejecting trials or data segments containing artifacts above a certain threshold would have resulted in the exclusion of the majority of data. ASR allowed us to keep most of the data while at the same time correcting for high-amplitude artifacts. We recommend to adjust the cut-off parameter to the specific dataset in order to find a good balance between artifact correction and signal retention (see Anders et al., 2020; Bloniasz, 2022)).

#### 4.2.7. Step 7: Interpolation of bad channels

Next, bad channels that were identified and removed before the ICA were interpolated using spherical splines. On average, 3.2 channels were removed for each dataset. This often included the vertical eye movement channel (EOG1), which was only accepted by around 10% of children and did not deliver usable signal for the remaining 90%. Since it was not relevant to our analyses, we did not interpolate the EOG1 channel.

#### 4.2.8. Step 8: Re-referencing

Finally, the EEG was re-referenced to the average of both mastoids. Re-referencing was done late in the pipeline to prevent artifacts in the reference electrodes taking over on all other electrodes. Generally, all types of references are possible here, including common average reference. However, when using layouts including few channels, as is often done in studies with young children (32 channels or less), the reference type should be carefully considered (Junghöfer et al., 1999).

#### 4.2.9. Step 9: Trial segmentation

While all previous steps were done using the EEGLAB toolbox (Delorme and Makeig, 2004), we transitioned to Fieldtrip (Oostenveld et al., 2011) for trial segmentation and following steps. Fieldtrip allows the segmentation of trials with unequal lengths, as is often the case in naturalistic and mobile EEG studies.

#### 4.2.10. Step 10: Visual and automatic rejection of artifacts

Particularly in mobile EEG data, long segments of data may be recorded between or outside of experimental trials. For this reason, we recommend performing additional artifact identification and rejection on segmented data that contains only epochs of interest. In the present pipeline, segmented EEG data were visually inspected to identify segments of the data containing remaining artifacts. These visually identified artifacts were then used in subsequent steps differently, depending on analysis goals: For analyses focusing on short sections (a couple of seconds) within the multiple-minute-long trials segmented before, we re-segmented trials to a fixed length in relation to one of the event triggers (e.g., 1 second before and 2 seconds after an event). From these sub-trials, all containing visual artifacts identified before or amplitudes exceeding ±200 µV in any channel were removed. On average, 0.72 sub-trials (range: 0-7) were rejected per dataset in this step, and 15 datasets (out of 303) were entirely excluded because they did not include any artifact-free trials. Mean number of artifact-free trials was 4.86 (range: 0-13). For analyses that focused on the entire trial, only the parts of the trials containing visual artifacts identified before were rejected. Which approach to choose depends strongly on analysis techniques, aims, and methodological considerations.

### 4.3. Comparison with Maryland Analysis of Developmental EEG (MADE) pipeline

In order to assess the effectiveness of our custom preprocessing pipeline for increasing the signal-to-noise ratio of developmental mobile EEG data, we compared the continuous data preprocessed with our pipeline (before trial segmentation) with continuous data preprocessed with the MADE pipeline (Debnath et al., 2020; without trial segmentation). Only data from the encoding session of the study and the full collected sample (*n* = 105) were used for comparison.

The continuous version of the MADE pipeline was used, with the same filter settings as applied for our custom pipeline. No anti-aliasing, down-sampling, deletion of outer channel layer, or baseline correction were performed, and data were re-referenced to the average of both mastoids. Twelve datasets were completely rejected and not preprocessed by the MADE pipeline.

As the two preprocessing pipelines were compared before any artifact rejection steps on epoched data, both retained the same number of segments for each dataset. However, after preprocessing with the custom pipeline, significantly fewer 1s-segments exceeded an ± 150 µV amplitude threshold (*M* = 127.72, *SD* = 236.89, *n* = 105) compared to the MADE pipeline (*M* = 668.81, *SD* = 467.81, *n* = 93; *t*(92) = -12.426, *p* < .001). On average, 8.48% (SD = 17.1) of the custom-preprocessed data, and 40.3% (SD = 24.48) of the MADE-preprocessed data still exceeded this threshold after preprocessing. Consequently, the data preprocessed with the custom pipeline contained overall fewer high-amplitude segments compared with the data preprocessed with the MADE pipeline.

Figure 3 shows the power spectral density as well as the time-frequency representations of the 10 example datasets, which are a subset of the encoding session dataset, after preprocessing with the custom and MADE preprocessing pipelines. Both pipelines retain the expected spectral characteristics of the EEG signal, including the decrease in power with increasing frequency.

**Figure 3.**
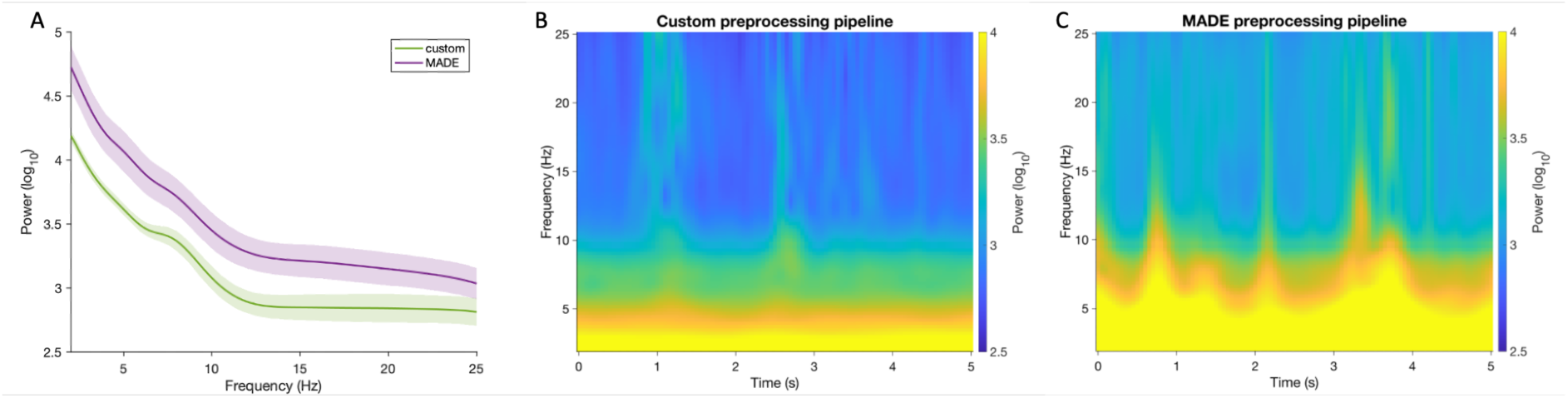
Comparison of spectral characteristics of the data after preprocessing with the custom and MADE preprocessing pipelines. (A) Mean power spectral density plot across the example datasets and all electrodes. (B) Mean time-frequency representations for the example datasets preprocessed with the custom pipeline, averaged across all electrodes. (C) Mean time-frequency representations for the example datasets preprocessed with the MADE pipeline, averaged across all electrodes. For the time-frequency decomposition, the first thirty 10s-segments of each example dataset were extracted. Time-frequency representations were then computed from the central 5s of each 10s-segment to minimize edge artifacts. We used complex Morlet wavelet convolution (width = 7) across frequencies from 2 Hz to 25 Hz in 0.25 Hz steps.

## 5. Discussion

In this article, we delineated the methodological challenges and practical solutions for collecting and preprocessing mobile EEG data from young children. In addition, we provide an example EEG dataset collected from freely moving toddlers during an episodic memory encoding task, demonstrating our preprocessing approach as well as providing measures of data quality.

The goal of this article is to facilitate the incorporation of mobile EEG in studies with young children. Mobile and naturalistic paradigms provide important opportunities to investigate cognitive development under real-life conditions. Adding EEG as a source of information to these paradigms enables researchers to investigate the neural processes underlying children’s active behavior and interactions with their environment. In the following, we focus on three major opportunities of mobile EEG for developmental cognitive neuroscience: (1) studying cognitive development in naturalistic environments; (2) including diverse populations that are difficult to study using stationary paradigms; and (3) investigating the interaction of neural activity with movement and goal-directed behavior.

### 5.1. Naturalistic environments

Mobile EEG is essential to study children’s neural processes and associated behaviors in their natural, real-world environments. This could include kindergartens and child-care centers (see Dopierala and Emberson, 2023), educational settings (e.g., Bevilacqua et al., 2019; Dikker et al., 2017; Dikker et al., 2020; Ko et al., 2017; Xu et al., 2022), children’s homes (see Dickinson et al., 2025; Troller-Renfree et al., 2021), public spaces (e.g., Deker and Pathman, 2021; Pathman et al., 2023 for behavioral examples), or group settings, allowing to study cognitive development in environments that are typically inaccessible to lab-based research. A crucial aspect of these naturalistic environments is that children usually do not inhabit them alone, but constantly engage with other people. These social interactions are essential for learning and exploration, and mobile EEG approaches offer the opportunity to investigate the neural processes underlying these interactions during naturalistic behavior. Existing parent-child brain-to-brain EEG (e.g., Liao et al., 2015; Norton et al., 2022; Turk et al., 2022) and hyperscanning approaches in classrooms (see Babiloni and Astolfi, 2014; Bevilacqua et al., 2019; Dikker et al., 2017; Janssen et al., 2021) could be extended to more naturalistic interaction settings and to other dyad or group types, including child-child interactions. Studying children in such diverse and social environments would extend and validate findings from controlled laboratory studies and open up new avenues for investigating how cognitive processes in childhood are shaped by real-world contexts.

### 5.2. Diverse populations

While most developmental neuroscience research relies on samples drawn from so-called WEIRD (Western, Educated, Industrialized, Rich, Democratic) populations, mobile EEG offers the possibility to increase diversity by increasing accessibility and reducing the constraints associated with lab-based recordings. Data collection in home or community environments could facilitate the inclusion of families from more diverse socioeconomic backgrounds and improve the representativeness of developmental EEG studies (see Dickinson et al., 2025; Janssen et al., 2021; Troller-Renfree et al., 2021). In addition, the portability of mobile EEG systems and reduced reliance on highly structured behavioral instructions may support cross-cultural research, allowing developmental processes to be studied in a wider range of cultural contexts (see Chin et al., 2025).

Mobile EEG may also enable the inclusion of developmental and neurodiverse populations that are typically difficult to study using stationary neuroimaging approaches. This includes young children (e.g., toddlers aged 1-3 years), as well as clinical populations such as children with attention deficit hyperactivity disorder (ADHD) or autism spectrum disorder (ASD), who may struggle with restricting their behavior and movement during stationary experiments. In these samples, the reduced movement constraints of mobile EEG studies may improve participation rates, participant motivation and well-being, and data quality, providing important insights into cognitive and neural processes during development in these populations (see Lau-Zhu et al., 2019).

### 5.3. Movement and goal-directed behavior

In early development, goal-directed behavior is often linked to bodily activity. Studying children’s neural processes while movement is taking place can provide important insights into brain-body couplings, embodied learning, and movement-cognition interactions in general (see Colomer et al., 2023; Lozada and Carro, 2016; Kontra et al., 2012; Laakso, 2011; Marmeleira and Duarte Santos, 2019; Pacheco et al., 2025). Behavioral research has already demonstrated that cognitive and sensorimotor development are intertwined (i.e., de Boer et al., 2026; Chung et al., 2024; Diamond, 2000; Lozada and Carro, 2016; Muentener et al., 2018; Pacheco et al., 2025; Schwarzer and Jovanovic, 2024) and physical changes in childhood relating to body size or proportions can affect cognitive processes such as perception or locomotor control (Adolph and Eppler, 1998; Adolph and Hoch, 2019; Franchak and Adolph, 2010; Witt, 2011). In adults, mobile brain/body imaging approaches integrate EEG recordings with motion tracking and other physiological recordings during active behavior to gain insights into neural processes associated with movement, and to use movement variables as an additional source of information in cognitive experiments (see Jungnickel et al., 2019; Makeig et al., 2009). Similar approaches could be pursued in children to better understand the interaction between movement, behavior, and cognition in early childhood. Examples are motor skill acquisition such as learning to walk or crawl, or sports-related movement and learning (see Contreras-Altamirano et al., 2025; Klapprott and Debener, 2026; Ludyga et al., 2016; Papin et al., 2024 for similar adult studies).

### 5.4. Open challenges and opportunities

Mobile EEG research faces a number of general challenges. A key issue is the trade-off between allowing free and naturalistic movement in unconstrained, ecologically valid settings while maintaining an acceptable signal-to-noise ratio and a certain degree of experimental control and replicability. A related challenge is to design mobile tasks that are comparable to stationary laboratory paradigms, which is important when trying to link and integrate findings across stationary and mobile EEG settings. These challenges are particularly pronounced when studying young children, who typically behave more spontaneously and less constrained in experimental settings compared to adults and may not consistently follow experimental instructions. At the same time, this could also be an advantage: Children’s behavior and EEG in these mobile paradigms is likely to more closely reflect real-world behavior and neural processes and be less influenced by the experimental context and social expectations than standard stationary assessments.

Given that the vast majority of mobile EEG studies are currently being conducted with adults, most currently available mobile EEG systems and methodological developments are designed and validated primarily for adult populations. Depending on children’s age, it may be difficult to find appropriately sized and suitable equipment, such as wearable solutions for mobile amplifiers and motion sensor suits, particularly when studying a range of age groups. Similarly, preprocessing pipelines and analysis approaches (e.g., the BeMoBIL pipeline, Klug et al., 2022; ICMoBI project, https://www.icmobi.org/) for mobile brain/body imaging research have been largely developed for adult data and require adaptation and validation across a range of childhood age ranges. Extending such approaches to child samples would offer the opportunity to move towards a more comprehensive investigation of brain, body, and behavior during early development.

Despite these challenges, mobile EEG also offers unique opportunities for developmental cognitive neuroscience. The design of experimental tasks and paradigms that align with children’s natural tendencies to play, explore, and move is likely to increase engagement, motivation, attention, and compliance, thereby enhancing the overall validity of results. In particular, allowing for spontaneous behavior in naturalistic contexts may facilitate the discovery of behaviors and related cognitive processes that are systematically overlooked by current, adult-designed experimental paradigms. In that sense, mobile EEG promotes a “bottom up” perspective into the study of child development.

## 6. Conclusions and Outlook

The present tutorial is meant to promote the adoption of mobile EEG in research on cognitive development in early childhood. By summarizing key methodological challenges and providing practical recommendations for data collection and preprocessing, alongside an example dataset and accompanying preprocessing script, we aim to lower entry barriers for researchers interested in adopting mobile EEG in their own work with young children.

The investigation of neural processes during active, naturalistic behavior holds potential for advancing our understanding of how children explore, learn, and develop in interaction with their own actions and their environment. Despite the methodological challenges associated with mobile EEG research in early and middle childhood, the approach described here demonstrates the feasibility of collecting and analyzing neural data from children during free and unconstrained movement.

## Funding sources

This work was supported by the Max Planck Institute for Human Development, Berlin, Germany, and the Max Planck School of Cognition, Leipzig, Germany, for AW.

## Acknowledgements

We are grateful to the participating families of our mobile EEG study for their time and commitment. We would also like to thank Christina Melnikov, Colin Oltrogge, Ece A. Sanin, Élena Reil, Helene Scheder-Bieschin, Jessica Schmidt, Kira J. Fahlbusch, Konstantin Kramer, Marie Springwald, Marius Fey, Nina Kolak, Saskia Taborski, Sebastian Rauch, Sophia Greiwe, Sophie Linzbach, Tallulah Axelrod, and Thalia Forbes-Mitchell for their invaluable help with data collection and Josefine Hild for her help in setting up the mobile EEG.

## Data availability

The example datasets and the preprocessing script are publicly available on: osf.io/x5tp8/

## Declaration of Generative AI and AI-assisted technologies in the writing process

During the preparation of this work the authors used ChatGPT in order to improve language and readability. After using this tool, the authors reviewed and edited the content as needed and take full responsibility for the content of the published article.

## Notes

https://www.osf.io/x5tp8/

### Competing Interest Statement

The authors have declared no competing interest.

